# Mechanism of *β*-Aminobutyric Acid-Induced Resistance in Wheat to the Grain Aphid, *Sitobion avenae* or Dissecting/Deciphering the Mechanism of *β*-Aminobutyric Acid-Induced Resistance in Wheat to the Grain Aphid, *Sitobion avenae*

**DOI:** 10.1101/001032

**Authors:** He-He Cao, Meng Zhang, Hui Zhao, Yi Zhang, Xing-Xing Wang, Shan-Shan Guo, Zhan-Feng Zhang, Tong-Xian Liu

## Abstract

The non-protein amino acid β-aminobutyric acid (BABA) could induce plant resistance to a broad spectrum of biotic and abiotic stresses. However, BABA-induced plant resistance to insects is less well-studied, especially its underlying mechanism. In this research, we applied BABA to wheat seedlings and tested its effects on *Sitobion avenae*. When applied as a soil drench, BABA significantly reduced weight of *S. avenae*, whereas foliar spray and seed treatment had no such effects. BABA-mediated suppression of *S. avenae* growth is dose dependent and could last at least for 7 days. The aminobutyric acid concentration in phloem sap of BABA-treated plants accumulated to high levels and increased with BABA concentrations applied. Moreover, after 10 days of treatment, the aminobutyric acid content in BABA-treated plants was still higher than that in control treatment. *S. avenae* could not discriminate artificial diet containing BABA from standard diet, indicating that BABA itself is not a deterrent to this aphid. Also *S. avenae* did not show preference for control plants or BABA-treated plants. Consistent with choice test results, *S. avenae* had similar feeding activities on control and BABA-treated plants, suggesting that BABA did not induce antifeedants in wheat seedlings. In addition, aminobutyric acid concentration in *S. avenae* feeding on BABA-treated plants was significantly higher than those feeding on control palnts. *S. avenae* growth rate was reduced on artificial diet containing BABA, indicating direct toxic effects of BABA to this aphid. These results suggest that BABA application could enhance wheat plant resistance to *S. avenae* and the mechanism is possibly due to direct toxicity of high BABA contents in plant phloem.

## Introduction

The non-protein amino acid β-aminobutyric acid (BABA) could enhance plant resistance against a broad spectrum of phytopathogens. BABA-induced plant resistance is effective against viruses, bacteria, oomycetes, fungi and phytopathogenic nematodes [1–4]. The mechanism of BABA-induced plant resistance to different pathogen is variable. BABA-induced *Arabidopsis* resistance against the oomycete pathogen *Peronospora parasitica* is based on callose deposition and independent of jasmonic acid (JA), salicylic acid (SA) and ethylene signaling pathways, whereas against the bacteria *Pseudomonas syringae* is solely dependent of SA and NPR1 [3]. Enhanced resistance of tobacco to tobacco mosaic virus by BABA application was found to be strictly dependent on SA pathway [2]. BABA-mediated grapevine resistance to downy mildew (*Plasmopara viticola*), however, is based on potentiating callose formation and JA signaling [5]. In addition to conferring plant resistance to pathogens, BABA also improves plant tolerance to salt, drought and high temperature, which is associated with accumulation of the abscisic acid [6–8]. *Arabidopsis* mutants that are insensitive to BABA-induced sterility have reduced resistance level to pathogen or tolerance to salt, suggesting that BABA-induced plant resistance to biotic and abiotic stresses has a genetic basis [9].

BABA also has been demonstrated to be effective in plant resistance to insects [10–12]. Hodge et al. (2005) found that applied BABA as a root drench to six legume plants reduced the performance of the pea aphid, *Acyrthosiphon pisum* (Harris) [10]. This is the first study examined the effects of BABA-mediated plant resistance to insects. When applied to Brassicaceae plants, BABA suppressed the growth of phloem-feeding insects *Myzus persicae* and *Brevicoryne brassicae* as well as chewing insects *Trichoplusia ni* and *Plutella xylostella* [11]. Recently, Tiwari et al. (2013) reported that BABA application could also induce citrus resistance to the Asian citrus psyllid, *Diaphorina citri* Ku wayama, and the authors did not find any direct toxicity of BABA to this insect by leaf-dipping bioassays [12]. Although these studies indicate that BABA could enhance plant resistance to insects, little is known about the underlying mechanism.

Plant direct resistance to insects relies largely on metabolites that exert toxic, antinutritive, or repellent effects. Insects could perceive these compounds by chemoreceptors and decide to accept or reject a host [13]. The phloem-feeding insects, like aphids and whiteflies, have evolved specialized mouthparts, the stylets, which penetrate plant epidermis, pass through intercellular tissue and the second wall material, searching for sieve element. During stylet penetration, aphids regularly puncture plant cells and taste cytosolic contents to decide to feed or leave before reaching phloem [14]. Once reached to the sieve element, aphids will always eject watery saliva to prevent sieve tube plugging before phloem sap ingestion. The probing behavior of aphids can be monitored by the electrical penetration graph (EPG) technique [15].

The English grain aphid, *Sitobion avenae* is a major pest of cereal worldwide [16]. This aphid causes substantial losses of cereal yield by removing photoassimilates and transmitting viruses. Application of chemical pesticides is still the main method to control this aphid; however, chemical control causes negative impacts on agroecosystem and can lead to insect resistance to pesticides [16]. Thus searching for alternative methods to control this aphid is of great significance.

Understanding the mechanism of BABA-induced plant resistance to insects will shed light on new methods of pest control. In this study we examined the effects of BABA and its isomers α-aminobutyric acid (AABA) and γ-aminobutyric acid (GABA) application to wheat on performance of the *S. avenae*. The durability of BABA-induced resistance was also investigated. Host preference and feeding behavior of *S. avenae* were studied to localized possible resistance factors. We found that BABA accumulated to high concentration in BABA-treated wheat phloem sap, so we use aphid artificial diet to test the direct toxicity of BABA to *S. avenae*.

## Materials and Methods

### Plants and Insects

Seeds of winter wheat, *Triticum aestivum* (var. XiNong 979), were germinated at room temperature (25 ± 1°C) for 2 days. Seedlings with similar size were then transplanted to pots (250 mL) containing a 5:1 mixture of peat moss (Pindstrup Mosebrug A/S, Ryomgaard, Denmark) and perlite. Seedlings were grown singly in pot unless otherwise indicated. Plants were cultivated in a walk-in growth chamber at 24 ± 1°C, 60 ± 5% relative humidity and a 14:10 hour light/dark regime. The plants were watered as necessary. The grain aphid, *S. avenae*, originally collected from a winter wheat field in Yangling, China, was reared on wheat seedlings (var. XiNong 979) in the same growth chamber.

### *S. avenae* Performance on Wheat Seedlings

Seven-day-old wheat seedlings were treated as indicated and then 2-3 apterous *S.avenae* adults were introduced to the first leaf of each seedling. The seedlings were caged by transparent plastic cages (8 cm in diameter and 30 cm in height) secured with nylon mesh at the top. One day later, the adults were removed, leaving 5-6 nymphs on the first leaf. After another 7 days, aphids on the first leaves were collected and weighed on a microbalance (resolution 0.001 mg; Sartorius MSA 3.6P-000-DM, Gottingen, Germany). To assess the effects of BABA treatment on plant growth, the fresh shoot weights of wheat seedlings were weighed after aphid collection. Weights of aphids feeding on wheat seedlings soil drenched with MilliQ water, different concentrations of BABA, and 50 mM GABA were analyzed using one-way analysis of variance (one-way ANOVA) following by Tukey’s HSD (Honestly Significant Difference) test. Mean aphids weights on plants soil drenched with 50 mM AABA, sprayed with BABA, or seed treatment with BABA were compared with respective control by Student’s *t*-test. Wheat seedlings weights were analyzed using one-way ANOVA following by Tukey’s HSD test. All statistical analyses in this research were conducted using IBM SPSS Statistics package (version 19.0; SPSS Inc., Chicago, IL, USA).

### *S. avenae* Settling Choice Tests

Seven-day-old wheat seedlings were soil drenched with 20 mL MilliQ water (control) or 25 mM BABA. One day after treatment, one control and one BABA-treated seedling were put in a transparent plastic cage (26 cm height × 15 cm length × 14 cm width) secured with nylon mesh at the top. Then 15 alate adults of *S. avenae* were introduced to each cage and the numbers of *S. avenae* on each seedling were recorded 4, 8, 24 and 48 h after the start of the experiment. This experiment was carried out in a completely dark room and 8 replicates were conducted. Numbers of aphids on control and BABA-treated plants at each time point were compared by paired *t*-test.

To determine whether BABA itself has direct antifeedant effects, we tested *S. avenae* preference to artificial diet containing BABA. The composition of standard artificial diet was based on Auclair (1965) [17] with modifications (Table S1). After sterile filtering through Millex GP syringe filters (hydrophilic polyethersulfone membrane, pore size 0.22 µm; Ireland), the artificial diets were stored at −70°C until use. The Petri dish lid (7 mm in height and 37 mm in diameter) was covered with a layer of stretched Parafilm^®^ M (Chicago, IL). Forty µL of MilliQ water (H_2_O), artificial diet (Control), and artificial diet containing 50 mM BABA were put on the Parafilm^®^ M separately. The test solutions were then covered with another layer of Parafilm^®^ M. We made a hole (5 mm in diameter) on the back of each lid and introduced twenty one 3nd-5th instar aphids through this hole. The numbers of aphids feeding on each test solution were recorded after 6, 12, 24, and 48 h. Fourteen replicates were performed. Percentages of aphids on each test solution within each recording time point were analyzed with one-way ANOVA and means were compared using Tukey’s HSD test.

### *S. avenae* Feeding Behavior

Aphid feeding activities on control and BABA-treated plant seedlings were recorded by the Giga-8 direct-current electrical penetration graph (DC-EPG) system [18]. The detail of this experiment can be found in our previous work [19]. Wheat seedlings were soil drenched with 20 mL MilliQ water (Control) or 25 mM BABA two days before EPG recording. Each apterous adult and wheat seedling was used once. Data were recorded by the Stylet+d software and analyzed with the Stylet+ software. The stylet pathway waveforms were distinguished according to Tjallingii (1978) [15]. EPG parameters were calculated using the Excel workbook for automatic parameter calculation of EPG data 4.3 [20]. Because the EPG data were not normally distributed, paired comparison of means of control treatment and BABA treatment was done by non-parametric Mann–Whitney *U*-test.

### Extraction and Analysis of Amino Acids

We used EDTA-facilitated exudation method to collect phloem sap of wheat seedling leaves for free amino acids analysis. Wheat seedlings were soil drenched as described 3 days before sample collection. The first leaf of each seedling was cut 1 cm above the leaf/stem junction and immediately put in 1.5 mL EP tube containing 600 µL EDTA solution (pH = 7.1). Two leaves were put in a tube and regarded as one replicate. Then these cut leaves were put in a completely dark growth chamber at 25°C 100% humidity for 3 hours. The samples were stored at −70°C until analysis.

Newly born *S. avenae* nymphs feeding on control and BABA treated plants for 7 days were collected and weighed. Free amino acids in aphids were extracted with 50% ethanol containing 0.1 M HCl. Mean concentrations of each amino acid from different treatments were compared by Student’s *t-*test.

The free amino acids extracted from plants and aphids were analyzed by LTQ XL™ linear ion trap mass spectrometer (Thermo Scientific, USA). Liquid chromatography separations were carried out with XTerra MS C18 Column (125Å pore size, 5 µm, 150 mm × 4.6 mm; Waters Corp., USA). Amino acids elution was performed applying a three-step gradient: A 100% for 7 min, 0-100% B linear for 2 min, 100% B for another 5 min, 0–100% A linear for 1 min, holding the system at 100% A for 5 min. Mobile phase A was a aqueous solution containing 5% acetonitrile and 0.1% formic acid; mobile phase B was 100% acetonitrile. The flow rate was 0.6 mL/min. The mass spectrometer worked in the positive electrospray ionization (ESI) mode. Nitrogen was used as the sheath gas (50.0 arbitrary units) and auxiliary gas (8.0 arbitrary units). The spray voltage was set at 4.5 kV and the ion transfer capillary temperature was 320°C. The amino acids were scanned and fragmented using data dependent MS/MS. Masses of precursor and product ions and collision energy for each amino acid were as described in Table S1. Data were acquired and processed using Xcalibur 2.1 software (Thermo Scientific). Quantification was achieved by external standard amino acid mixture of known concentrations (AA-S-18, Sigma). For our method can not detect glycine, our data do not include glycine.

### Time Course Assay

To investigate the durability of the effects of BABA treatment on *S. avenae* performance and phloem sap amino acids composition, we conducted a time course experiment. Seven-day-old wheat seedlings were soil drenched with 20 mL MilliQ water or 25 mM BABA, and aphid performance assays started 0, 7, and 14 days after treatment; phloem sap were collected 3, 10, and 17 days after treatment for amino acids analysis. Aphid weights and aminobutyric acid concentrations of different treatments within each sample day were analyzed by Student’s *t*-test respectively.

### Peroxidase Assays

Peroxidase (POD) is involved in some forms of plant resistance to insects [21]. The first leaves of control and BABA-treated plants were collected 2, 4, and 6 days after treatment, extracted and analyzed as described previously [19]. POD activities of control and BABA-treated plants within each sample day were compared by Student’s *t*-test. In another experiment, plants were divided to following four treatments: (1) control plants (soil drenched with water); (2) control plants infested with fifteen 3rd instar-adult *S. avenae*; (3) BABA-treated plants (soil drenched with 25 mM BABA); (4) BABA-treated plants infested with fifteen 3rd-adult *S. avenae*. After 2, and 4 days of treatment, the first leaves were collected and analyzed as described. Mean activities of POD among treatments on each day were analyzed by one-way ANOVA, with Tukey’s HSD post-hoc test.

### Artificial Diet Assays

We tested the direct impacts of BABA on performance of *S. avenae* by measuring *S. avenae* nymph growth on standard artificial diet and artificial diet containing 50 mM BABA, AABA, or GABA. We use AABA and GABA in artificial diet to exclude the possibility that the imbalance of artificial diet have negative impacts on aphid growth. Thirty-five µL of artificial diet was confined between two layers of stretched Parafilm^®^ M on plastic cylinder (1 cm in height and 1cm in diameter) and the artificial diet was replaced every two days. Five one-day-old nymphs produced by apterous *S. avenae* were transferred to one tube containing one kind of artificial diets and regarded as one replicate. The weight of individual nymph was weighed on the MSA 3.6P-000-DM microbalance after feeding on artificial diet for 4 days. We also tested aphid performance on artificial diet containing 50 mM and 100 mM BABA with similar manner, but the one day old nymphs used were produced by *S. avenae* collected from wheat seedlings (var. XiNong 979) in greenhouse. Weights of aphids were analyzed with one-way ANOVA following by Tukey’s HSD test.

## Results

### *S. avenae* Performance

Compared with control treatment, BABA applied as a soil drench significantly reduced weights of *S. avenae* on wheat seedlings and the effects increased with BABA concentration (Fig. 1A). In contrast, soil drench with GABA (Fig. 1A) and AABA (Fig. 1B), spray with BABA (Fig. 1C), or seed treatment with BABA (Fig. 1D) had no negative impacts on *S. avenae* weights.

**Figure 1.**
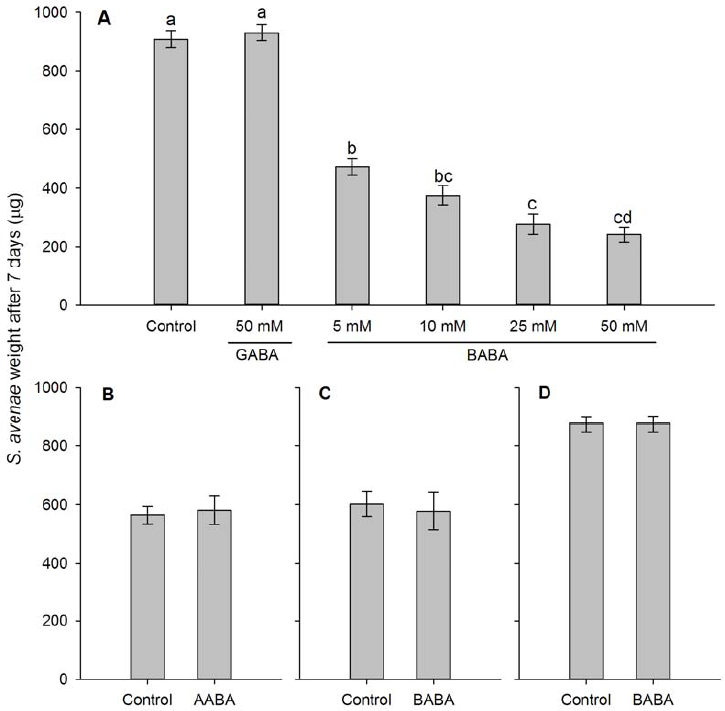
*Sitobion avenae* weights after feeding for 7 days on wheat seedlings with different treatments. Aphid weights on wheat plants (A) soil drenched with MilliQ water (control), different concentrations of BABA, 50 mM GABA, (B) soil drenched with MilliQ water and 50 mM AABA, (C) sprayed with MilliQ water and 50 mM BABA, (D) seed treatment with water and 50 mM BABA. Bars represent mean ± SEM. Different letters above bars in fig. 1A indicate statistically significant differences (*P* < 0.05, Turkey’s HSD test).

### *S. avenae* Preference

BABA-treated wheat plants were not attractive or repellent to *S. avenae* (Fig. 2A). *S. avenae* preferred artificial diets than MilliQ water, but showed no preference between standard artificial diet and artificial diet containing 50 mM BABA (Fig. 2B).

**Figure 2.**
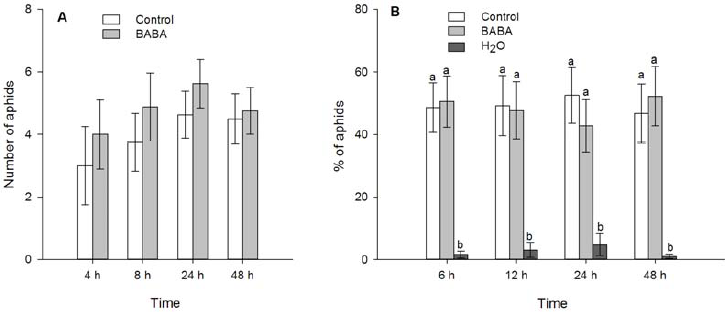
Settling preference of *Sitobion avenae*. (A) Mean number (± SEM) of *S. avenae* settling on control or BABA-treated plants (n = 8, paired *t*-test). Plants were soil drenched with MilliQ water (control) or 25 mM BABA. (B) Percentage of *S. avenae* on MilliQ water (H_2_O), standard artificial diet (control), and artificial diet containing 50 mM BABA (BABA). Shown are mean ± SEM (n = 14). Different letters indicate statistically significant differences (*P* < 0.05, Turkey’s HSD test).

### Free Amino Acids Composition of Phloem Sap

The relative concentration of aminobutyric acid in phloem sap of BABA-treated wheat plants was higher than that in control plants (Fig. 3). Our method can not discriminate BABA from its isomers, the excess amount of aminobutyric acid in BABA-treated plants was assumed to be BABA. Because wheat plants could produce GABA not BABA, we consider the aminobutyric acid in control plants was GABA. Soil drenched with AABA did not influence aminobutyric acids concentration in wheat phloem, while GABA only slightly increased aminobutyric acids levels.

**Figure 3.**
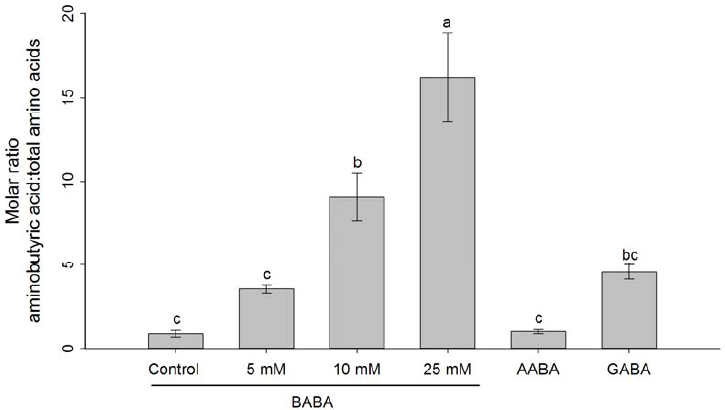
Relative aminobutyric acid levels in phloem sap of wheat seedlings. Plants were soil drenched with different concentrations of BABA, 50 mM AABA and 50 mM GABA. Phloem sap was collected 3 days after treatment. Shown are mean ± SEM (n = 6). Different letters indicate statistically significant differences (*P* < 0.05, Turkey’s HSD test).

Except BABA, the relative concentration of serine in BABA-treated wheat phloem sap was also significantly less than that in control plants (Student’s *t*-test, *t* = 2.78, df = 10, *P* = 0.02; Fig. S2), while other amino acids levels between treatments were similar (Fig. S2).

### Free Amino Acids Composition of *S. avenae*

The relative concentration of aminobutyric acid in *S. avenae* feeding on BABA-treated plants was significantly higher than those feeding on control plants (Student’s *t*-test, *t* = 8.84, df = 6, *P* < 0.001; Fig. 4). The high levels aminobutyric acid in *S. avenae* should be BABA from phloem sap of BABA-treated plants. The relative levels of alanine (*P* = 0.052) and threonine (*P* = 0.073) in aphids feeding on BABA-treated plants were slightly lower than those feeding on control plants (Fig. 4). The other amino acids composition pattern was similar.

**Figure 4.**
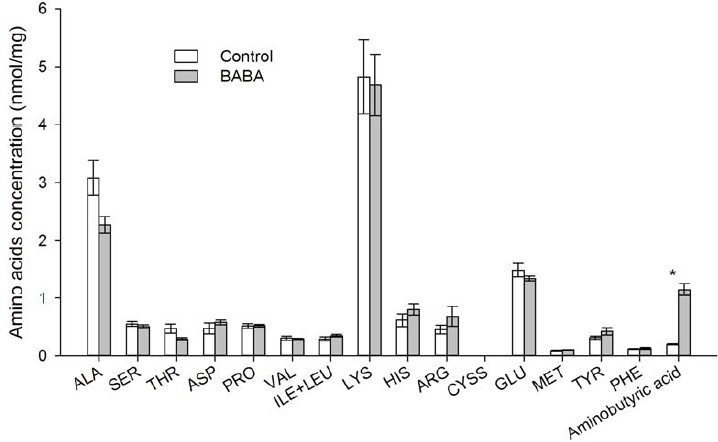
Amino acids concentration in *Sitobion avenae* bodies. Wheat plants were soil drenched with MilliQ water (control) and 25 mM BABA. Newly born nymphs feeding on control or BABA-treated plants for 7 days were collected for analysis. Shown are mean ± SEM (n = 4, \**P* < 0.05, Student’s *t*-test).

### Durability of BABA Treatment

After 7 days of BABA treatment, wheat seedlings still could reduce *S. avenae* growth (Student’s *t*-test, *t* = 3.10, df = 56, *P* < 0.01; Fig. 5A), which is positively correlated with aminobutyric acid levels in plant phloem sap (Fig. 5B). The relative aminobutyric acid concentration in BABA-treated plants was higher than that in control plants 3 days and 10 days after treatment, but not significant after 17 days (Fig. 5B).

**Figure 5.**
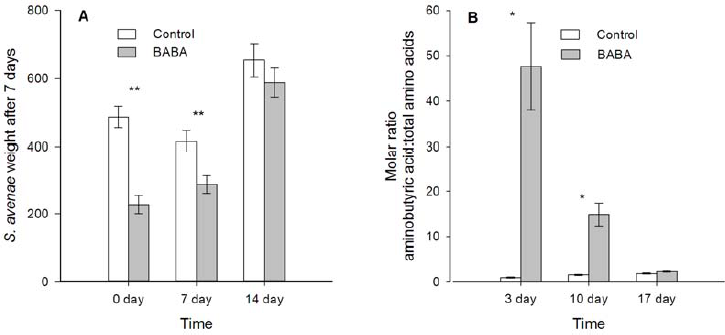
Durability of BABA treatment on *Sitibion avenae* performance and relative aminobutyric content in plant phloem. (A) Weights of *S. avenae* after feeding on control and BABA-treated plants for 7 days. Performance assay started 0, 7, and 14 day after treatment. (B) Relative aminobutyric acid levels in wheat seedlings phloem, which was collected 3, 10, and 17 days after treatment. Shown are mean ± SEM (\**P* < 0.05, \*\**P* < 0.01, Student’s *t*-test).

### Feeding Activities of *S. avenae*

The feeding behavior of *S. avenae* on BABA-treated and control plants was similar revealed by EPG technique (Table 1). The number of probes, total duration of probing, and total duration of phloem feeding (E2) were not significantly different, suggesting that BABA treatment did not induce physical barriers or chemical deterrents in wheat seedlings against *S. avenae* (Table 1).

**Table 1.** EPG parameters of *Sitobion avenae* during an 8 h recording on wheat seedlings soil drenched with water (Control) or beta-aminobutyric acid (BABA) 48 h earlier.

| EPG Parameters | Control<br>n=21 | BABA<br>n=25 |
| --- | --- | --- |
| Number of probes | 10.7 ± 1.1 | 13.9 ± 2.1 |
| Number of short probes (<3 min) | 4.0 ± 0.6 | 5.5 ± 1.2 |
| Number of probes to the 1st E1 | 7.8 ± 1.3 | 8.6 ± 1.0 |
| Time from 1st probe to 1st E1 | 2.3 ± 0.4 | 2.6 ± 0.3 |
| Total duration of probing (h) | 7.1 ± 0.1 | 7.0 ± 0.2 |
| Total duration of pathway | 2.1 ± 0.2 | 2.3 ± 0.2 |
| Total duration of E2 (h) | 4.1 ± 0.3 | 3.8 ± 0.4 |
| Mean duration of E2 (h) | 1.7 ± 0.3 | 2.4 ± 0.5 |
| Total duration of E1 (min) | 16.0 ± 4.1 | 12.8 ± 2.3 |
| Mean duration of E1 (min) | 5.9 ± 1.6 | 5.2 ± 1.2 |
| Number of E2 | 3.0 ± 0.3 | 2.6 ± 0.4 |
| Number of E1 | 3.1 ± 0.4 | 2.9 ± 0.4 |
| Duration of xylem ingestion (h) | 0.6 ± 0.2 | 0.6 ± 0.1 |

Values represent means ( ± SEM). E1: Salivation phase; E2: Phloem sap ingestion. n: number of replicates.

### POD Activities

There were no difference of POD activities between control and BABA-treated plants (Fig. S3A). Also BABA treatment did not increase POD activities of aphid infested wheat seedlings compared with aphid infested control plants (Fig. S3B).

### Direct Toxicity of BABA on *S. avenae*

*S. avenae* feeding for 4 days on artificial diets containing BABA or GABA had lower weight than those feeding on standard artificial diet; whereas those feeding on artificial diet containing AABA had no influence on aphid weight (one-way ANOVA, *F* = 11.92, df = 3, 35, *P* < 0.001; Fig. 6A). *S. avenae* feeding on artificial diet containing BABA had only 55% weight of those feeding on standard artificial diet (Fig. 6A). *S. avenae* feeding for 3 days on standard artificial diet had higher weight than those feeding on artificial diets containing 50 mM or 100 mM BABA (one-way ANOVA, *F* = 11.33, df = 2, 49, *P* < 0.001; Fig. 6B). The mortality of *S. avenae* feeding on all artificial diets was similar (data not shown).

**Figure 6.**
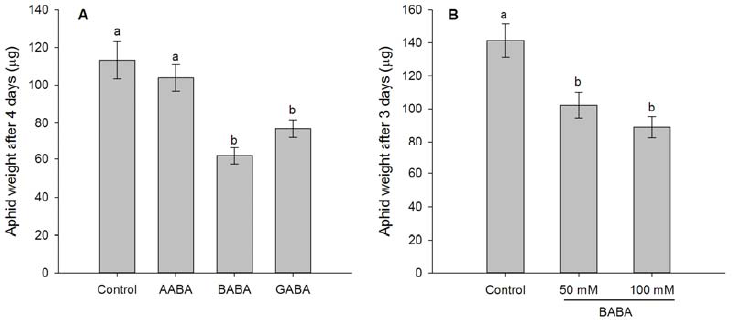
Direct toxicity of BABA to *Sitobion avenae*. (A) Weights of nymphs after feeding on standard artificial diet (control) and artificial diet containing 50 mM AABA, BABA or GABA for 4 days. (B) Weights of nymphs after feeding on standard artificial diet (control) and artificial diet containing 50 mM BABA or 100 mM BABA for 3 days. Shown are mean ± SEM. Different letters indicate statistically significant differences (*P* < 0.05, Turkey’s HSD test).

## Discussion

Although there have been numerous studies examined the effects and mechanisms of BABA-mediated plant resistance to plant pathogens, few papers investigated the role of BABA in plant resistance to insects [4,9]. The already published papers suggest that BABA application could suppress performance of some aphids, caterpillars, and psyllid [10–12]. Our results demonstrated that BABA applied as a root drench could significantly reduce weight of *S. avenae* and the effects was dose dependent. We found high concentration of BABA in treated plant phloem sap, and the BABA level was still higher than that in control phloem sap after 10 days, suggesting that wheat could absorb BABA by roots and metabolize or decompose BABA slowly. The BABA-mediated suppression of aphid growth could last at least for 7 days, which is correlated with BABA contents in phloem sap. However, we did not find any influence of BABA application on *S. avenae* host preference or feeding behavior, indicating that BABA did not induce any repellents in wheat seedlings. It was assumed that resistance to insects induced by BABA is not based on its direct toxicity [10,12]. In contrast, we found that *S. avenae* feeding on artificial diet containing BABA had reduced weight, suggesting that BABA has direct toxic effects on *S. avenae*.

Our results demonstrated that only when drenched into the soil, BABA could reduce *S. avenae* growth on wheat. And BABA-mediated wheat resistance to *S. avenae* is positively correlated with BABA concentration in wheat phloem sap. By using ^14^C-labled BABA, Cohen and Gisi (1994) found that only a small proportion of sprayed BABA is taken up by plant leaves [22]. Foliar spray and seed treatment with BABA may result in relative lower BABA concentration in wheat phloem and thus have no effects on aphid performance. Root drench with BABA rendered wheat plants a durable resistance to *S. avenae*, which corresponds to high BABA concentration in wheat phloem. Our time course experiment showed that BABA concentration in treated plants decreased with time, which suggests that BABA could be metabolized by wheat plants slowly.

*S. avenae* cannot discriminate artificial diet containing BABA from standard artificial diet, implying that *S. avenae* may not have corresponding receptors. Because BABA is rarely found in plants, aphids have few chances to encounter this compound in nature and do not evolve the ability to perceive BABA [23]. The EPG data indicate that *S. avenae* exhibited similar feeding activities on control and BABA-treated plants, which is consistent with host preference results. BABA-induced plant resistance to pathogens is mainly through callose formation and SA, ABA signaling pathway [3,9,24]. BABA induced callose deposition at sits of pathogen penetration; thus preventing spread of pathogen [3,24]. Callose is also involved in plant phloem sealing mechanisms, which could confer plant resistance to aphids [25,26]. Aphids inject watery saliva to prevent plugging and sealing of sieve plates [26]. This ejection of saliva could be detected by the EPG as E1 salivation. However, total or mean duration of E1 on control and BABA-treated plants was not significantly different. And aphids had similar total as well as mean duration of phloem sap ingestion. Therefore, BABA-mediated wheat resistance to *S. avenae* is not possibly based on callose induced phloem occlusion. JA is the most important cellular signal in plant immunity to most insect herbivores [27]. Application of JA and its derivatives methyl jasmonate could increase plant secondary metabolites and resistance to a broad spectrum of insects [27,28]. Our previous work showed that both methyl jasmonate and SA treatment deterred *S. avenae* preference and feeding activities, but had no effects on aphid performance [19]. These rule out the possible involvement of JA and SA signaling transduction pathways in the BABA-mediated wheat resistance to *S. avenae*.

It has been believed that BABA-mediated plant resistance to pathogens and insects is not based on its direct toxicity [10,12,23]. Recently, however, Šašek et al. (2012) found BABA had direct antifungal activity against *Leptosphaeria maculans* and the effect was comparable with the fungicide tebuconazole [29]. Our study showed that both BABA and GABA had direct toxicity to *S. avenae*, when added to artificial diet. GABA is the major inhibitory neurotransmitter in the vertebrates and invertebrates nervous system [30,31]. Synthetic diet containing GABA reduced growth and survival of oblique-banded leaf roller larvae [32]. GABA possibly reduced *S. avenae* performance by disturbing aphid nervous system [31]. However, the mechanism of BABA direct suppression of aphid growth is still unknown. In contrast to our findings, Hodge et al. (2005) did not find direct toxicity of BABA to the pea aphid *Acyrthosiphon pisum* and Tiwari (2013) reported that BABA had no influence on Asian citrus psyllid survival [10,12]. It is possible that their application methods only lead to low accumulation of BABA in insects, which is not high enough to reduce insects’ fitness. Our data showed that the aminobutyric acid concentration in *S. avenae* feeding on wheat plant treated with BABA was 5.8 times higher than that on control plants. The excess aminobutyric acid in *S. avenae* is assumed to be BABA. Another possibility is that the direct toxicity of BABA to insects is species-specific.

BABA treatment could alter amino acids balance in *Arabidopsis*, and L-glutamine could inhibit BABA-induced resistance to thermotolerance and a bacterial pathogen [33,34]. We found that BABA-treated wheat phloem accumulated lower serine than water-treated control. These findings imply that BABA possibly involves in plant amino acids metabolite. *S. avenae* feeding on BABA treated wheat seedlings had lower alanine and threonine in their bodies. The lower alanine concentration in *S. avenae* feeding on BABA-treated plants may not be due to alanine difference in plant phloem, because BABA-treated wheat had more alanine. Therefore, this suggests that BABA could also change aphid amino acids metabolite by unknowing mechanism. Furthermore, glycine is an important inhibitory neurotransmitter in the central nervous system, while BABA is a partial agonist at the glycine receptor [35]. Low concentrations of BABA competitively inhibits glycine responses, whereas higher concentrations elicits a significant membrane current [35]. Therefore, BABA possibly exerts its direct effects to *S. avnae* by causing aphid neurological disorders.

BABA treatment did not influence aphid host preference and feeding activities, but resulted in lowered aphid weight and durable high concentration of BABA in wheat phloem sap. Using artificial diet, we clearly showed that BABA had direct toxic effects to *S. avenae*. These findings expand our knowledge of BABA-mediated plant resistance to insects. Further research is needed to investigate whether BABA has direct toxicity to other insects as well as non-target organisms.

## Acknowledgments

### Author Contributions

Conceived and designed the experiments: HHC TXL. Performed the experiments: HHC HZ ZFZ XXW. Analyzed the data: HHC MZ ZFZ. Contributed reagents/materials/analysis tools: ZM YZ XXW SSG. Wrote the paper: HHC TXL.

## References

1. Oka Y, Cohen Y, Spiegel Y (1999) Local and systemic induced resistance to the root-knot nematode in tomato by DL-*β*-amino-n-butyric acid. Phytopathology 89: 1138–1143.

2. Siegrist J, Orober M, Buchenauer H (2000) *β*-Aminobutyric acid-mediated enhancement of resistance in tobacco to tobacco mosaic virus depends on the accumulation of salicylic acid. Physiol Mol Plant Pathol 56: 95–106.

3. Zimmerli L, Jakab G, Metraux JP, Mauch-Mani B (2000) Potentiation of pathogen-specific defense mechanisms in *Arabidopsis* by *β*-aminobutyric acid. Proc Natl Acad Sci U S A 97: 12920–12925.

4. Justyna P-G, Ewa K (2013) Induction of resistance against pathogens by *β*-aminobutyric acid. Acta Physiologiae Plantarum 35: 1735–1748.

5. Hamiduzzaman MM, Jakab G, Barnavon L, Neuhaus J-M, Mauch-Mani B (2005) *β*-Aminobutyric acid-induced resistance against downy mildew in grapevine acts through the potentiation of callose formation and jasmonic acid signaling. Mol Plant-Microbe Interact 18: 819–829.

6. Jakab G, Ton J, Flors V, Zimmerli L, Metraux JP, et al. (2005) Enhancing Arabidopsis salt and drought stress tolerance by chemical priming for its abscisic acid responses. Plant Physiol 139: 267–274.

7. Zimmerli L, Hou B-H, Tsai C-H, Jakab G, Mauch-Mani B, et al. (2008) The xenobiotic *β*-aminobutyric acid enhances *Arabidopsis* thermotolerance. The Plant Journal 53: 144–156.

8. Du Y-L, Wang Z-Y, Fan J-W, Turner NC, Wang T, et al. (2012) *β*-Aminobutyric acid increases abscisic acid accumulation and desiccation tolerance and decreases water use but fails to improve grain yield in two spring wheat cultivars under soil drying. J Exp Bot 63: 4849–4860.

9. Ton J, Jakab G, Toquin V, Flors V, Iavicoli A, et al. (2005) Dissecting the beta-aminobutyric acid-induced priming phenomenon in *Arabidopsis*. Plant Cell 17: 987–999.

10. Hodge S, Thompson GA, Powell G (2005) Application of DL-*β*-aminobutyric acid (BABA) as a root drench to legumes inhibits the growth and reproduction of the pea aphid *Acyrthosiphon pisum* (Hemiptera : Aphididae). Bull Entomol Res 95: 449–455.

11. Hodge S, Pope TW, Holaschke M, Powell G (2006) The effect of *β*-aminobutyric acid on the growth of herbivorous insects feeding on Brassicaceae. Ann Appl Biol 148: 223–229.

12. Tiwari S, Meyer WL, Stelinski LL (2013) Induced resistance against the Asian citrus psyllid, *Diaphorina citri*, by *β*-aminobutyric acid in citrus. Bull Entomol Res 103: 592–600.

13. Mithofer A, Boland W (2012) Plant defense against herbivores: chemical aspects. Annu Rev Plant Biol 63: 431–450.

14. Powell G, Tosh CR, Hardie J (2006) Host plant selection by aphids: behavioral, evolutionary, and applied perspectives. Annu Rev Entomol 51: 309–330.

15. Tjallingii WF (1978) Electronic recording of penetration behaviour by aphids. Entomol Exp Appl 24: 721–730.

16. Emden HFv, Harrington R (2007) Aphids as Crop Pests. CABI, Wallingford, United Kingdom: 1–717.

17. Auclair JL (1965) Feeding and nutrition of the pea aphid, *Acyrthosiphon pisum* (Homoptera: Aphidae), on chemically defined diets of various pH and nutrient levels. Ann Entomol Soc Am 58: 855–875.

18. Prado E, Tjallingii WF (1994) Aphid activities during sieve element punctures. Entomol Exp Appl 72: 157–165.

19. Cao H-H, Wang S-H, Liu T-X (2013) Jasmonate and salicylate induced defenses in wheat affect host preference and probing behavior but not performance of the grain aphid, *Sitobion avenae*. Insect Sci: (in press).

20. Sarria E, Cid M, Garzo E, Fereres A (2009) Excel Workbook for automatic parameter calculation of EPG data. Comput Electron Agric 67: 35–42.

21. Liu X, Williams CE, Nemacheck JA, Wang H, Subramanyam S, et al. (2010) Reactive oxygen species are involved in plant defense against a gall midge. Plant Physiol 152: 985–999.

22. Cohen Y, Gisi U (1994) Systemic translocation of ^14^C-DL-3-aminobutyric acid in tomato plants in relation to induced resistance against *Phytophthora infestans*. Physiol Mol Plant Pathol 45: 441–456.

23. Jakab G, Cottier V, Toquin V, Rigoli G, Zimmerli L, et al. (2001) *β*-Aminobutyric acid-induced resistance in plants. Eur J Plant Pathol 107: 29–37.

24. Ton J, Mauch-Mani B (2004) *β*-Amino-butyric acid-induced resistance against necrotrophic pathogens is based on ABA-dependent priming for callose. The Plant Journal 38: 119–130.

25. Will T, van Bel AJE (2006) Physical and chemical interactions between aphids and plants. J Exp Bot 57: 729–737.

26. Tjallingii WF (2006) Salivary secretions by aphids interacting with proteins of phloem wound responses. J Exp Bot 57: 739–745.

27. Howe GA, Jander G (2008) Plant immunity to insect herbivores. Annu Rev Plant Biol 59: 41–66.

28. Zhang H, Memelink J (2009) Regulation of secondary metabolism by jasmonate hormones. In: Osbourn AE, Lanzotti V, editors. Plant-derived Natural Products: Springer US. pp. 181–194.

29. Šašek V, Nováková M, Dobrev PI, Valentová O, Burketová L (2012) *β*-Aminobutyric acid protects *Brassica napus* plants from infection by *Leptosphaeria maculans*. Resistance induction or a direct antifungal effect? Eur J Plant Pathol 133: 279–289.

30. Huebner CA, Holthoff K (2013) Anion transport and GABA signaling. Frontiers in Cellular Neuroscience 7.

31. Hosie A, Sattelle D, Aronstein K, ffrench-Constant R (1997) Molecular biology of insect neuronal GABA receptors. Trends Neurosci 20: 578–583.

32. Ramputh A-I, Bown AW (1996) Rapid *γ*-aminobutyric acid synthesis and the inhibition of the growth and development of oblique-banded leaf-roller larvae. Plant Physiol 111: 1349–1352.

33. Singh PK, Wu C-C, Zimmerli L (2010) Beta-aminobutyric acid priming by stress imprinting. Plant Signaling & Behavior 5: 878–880.

34. Wu CC, Singh P, Chen MC, Zimmerli L (2010) L-Glutamine inhibits beta-aminobutyric acid-induced stress resistance and priming in *Arabidopsis*. J Exp Bot 61: 995–1002.

35. Schmieden V, Betz H (1995) Pharmacology of the inhibitory glycine receptor: agonist and antagonist actions of amino acids and piperidine carboxylic acid compounds. Mol Pharmacol 48: 919–927.

